# Unsupervised explainable AI for the collective analysis of a massive number of genome sequences: various examples from the small genome of pandemic SARS-CoV-2 to the human genome

**DOI:** 10.1101/2021.05.23.445371

**Authors:** Toshimichi Ikemura, Yuki Iwasaki, Kennosuke Wada, Yoshiko Wada, Takashi Abe

## Abstract

In genetics and related fields, huge amounts of data, such as genome sequences, are accumulating, and the use of artificial intelligence (AI) suitable for big data analysis has become increasingly important. Unsupervised AI that can reveal novel knowledge from big data without prior knowledge or particular models is highly desirable for analyses of genome sequences, particularly for obtaining unexpected insights. We have developed a batch-learning self-organizing map (BLSOM) for oligonucleotide compositions that can reveal various novel genome characteristics. Here, we explain the data mining by the BLSOM: unsupervised and explainable AI. As a specific target, we first selected SARS-CoV-2 (severe acute respiratory syndrome coronavirus 2) because a large number of the viral genome sequences have been accumulated via worldwide efforts. We analyzed more than 0.6 million sequences collected primarily in the first year of the pandemic. BLSOMs for short oligonucleotides (e.g., 4~6-mers) allowed separation into known clades, but longer oligonucleotides further increased the separation ability and revealed subgrouping within known clades. In the case of 15-mers, there is mostly one copy in the genome; thus, 15-mers appeared after the epidemic start could be connected to mutations. Because BLSOM is an explainable AI, BLSOM for 15-mers revealed the mutations that contributed to separation into known clades and their subgroups. After introducing the detailed methodological strategies, we explained BLSOMs for various topics. The tetranucleotide BLSOM for over 5 million 5-kb fragment sequences derived from almost all microorganisms currently available and its use in metagenome studies. We also explained BLSOMs for various eukaryotes, such as fishes, frogs and *Drosophila* species, and found a high separation ability among closely related species. When analyzing the human genome, we found evident enrichments in transcription factor-binding sequences (TFBSs) in centromeric and pericentromeric heterochromatin regions. The tDNAs (tRNA genes) were separated by the corresponding amino acid.

## INTRODUCTION

The Kohonen self-organizing map (SOM), an unsupervised neural network algorithm, is a powerful tool for the clustering and visualization of high-dimensional complex data on a two-dimensional map (Kohonen et al., 1996). We modified the conventional SOM for genome informatics based on batch learning to make the learning process and the resulting map independent of the order of data input (Kanaya et al., 2001). The batch-learning SOM (BLSOM) is suitable for high-performance parallel computing and thus for big data analysis (Abe et al., 2003, 2006a). It is also worth mentioning that we can obtain unexpected insights by daring to leave knowledge discovery to AI. By analyzing the compositions of short oligonucleotides (e.g., 4- and 5-mers) in a large number of genomic fragments (e.g., 10 kb) derived from a wide variety of species, the BLSOM enables the separation (self-organization) of the genomic sequences by species and phylogeny and identifies oligonucleotides that significantly contribute to the separation (Abe et al., 2003, 2006c; Iwasaki et al., 2013b; Bai et al., 2014; Kikuchi et al., 2015). In an analysis of the genomic fragments of a wide range of microbial genomes, over 5 million sequences were successfully separated by phylogenetic groups with high accuracy (Abe et al., 2020). A BLSOM program suitable for PC cluster systems is available on the following website: http://bioinfo.ie.niigata-u.ac.jp/?BLSOM.

In our previous analysis of all influenza A strains, the viral genomes were separated (self-organized) by host animals based only on the similarity of the oligonucleotide composition, even though no host information was provided during the machine learning process (Iwasaki et al., 2011a, b): unsupervised AI. Notably, the BLSOM is also an explainable AI that can identify diagnostic oligonucleotides, which contribute to host-dependent clustering (self-organization). When studying the 2009 swine-derived flu pandemic (H1N1/2009), we detected directional time-series changes in the oligonucleotide composition due to possible adaptations to the new host, namely, humans (Iwasaki et al., 2011a), and these findings have shown that near-future prediction was possible, albeit partially (Iwasaki et al., 2013a). We found similar time-series changes in the oligonucleotide composition for ebolavirus, MERS coronavirus (Wada et al., 2016, 2017) and SARS-CoV-2 (Wada et al., 2020b; Ikemura et al., 2020; Iwasaki et al., 2021; Abe et al., 2021). In the present paper, SARS-CoV-2 genomes, which are of great interest to society, were used as an operative example to illustrate how this unsupervised explainable AI supports efficient knowledge discovery from big sequence data.

### Analyses of SARS-CoV-2

To confront the global threat of COVID-19, a massive number of SARS-CoV-2 genome sequences have been decoded and published by GISAID (Elbe and Buckland-Merret, 2017; https://www.gisaid.org/). SARS-CoV-2 is an RNA virus with a rapid evolutionary rate, and based on variant types, eight clades have already been defined by GISAID, but the diversity may be far greater. Due to the current explosive increase in available sequences, we must develop new technologies that can grasp the whole picture of big-sequence data and support efficient data mining. We have recently analyzed time-series changes in short and long oligonucleotide compositions in a large number of SARS-CoV-2 genomes and found many oligonucleotides that are expanding rapidly in the virus population, which allowed us to predict candidate advantageous mutations for growth in human cells (Wada et al., 2020b; Ikemura et al., 2020). Furthermore, the oligonucleotide BLSOM can classify the virus sequences into not only the known clades but also their subgroups (Abe et al., 2021). After the above-mentioned publications, a large number of sequences of this virus have continued to accumulate and will exceed one million. These big data, which have been acquired through worldwide efforts, are valuable not only for dealing with the current and future epidemics of infectious viruses but also for basic research in genetics and related fields. In this paper, we first used SARS-CoV-2 sequences, including the new sequences, to explain how unsupervised explainable AI enables knowledge discovery from a massive number of genome sequences.

### BLSOM of short oligonucleotides

In this study, approximately 0.7 million SARS-CoV-2 genomes, which were isolated from December 2019 to February 2021, were analyzed, and the polyA-tail was removed prior to the analysis. Figure 1 shows BLSOMs for di- to hexanucleotide compositions in all viral genomes; importantly, no information other than the oligonucleotide composition was provided during the machine learning step (unsupervised AI). The total number of grid points (i.e., nodes) was set to 1/100 of the total number of viral genomes. After learning, to determine whether the separation achieved with the BLSOM was related to known clades published by GISAID, grid points containing genomes of a single clade are colored to indicate each clade, and grid points containing those of multiple clades are displayed in black. As shown in Fig. 1A, a major portion of grid points on the dinucleotide BLSOM are black, but a major portion of the grids on penta- and hexanucleotide BLSOMs are colored, showing the good classification power of the BLSOMs (Figs. 1A and 1Bi). Notably, the grid point where even one other clade-derived sequence is mixed is displayed in black, despite of the condition that an average of 100 sequences per grid point was set. This condition may be very strict; thus, in the hexanucleotide BLSOM shown in Fig. 1Bii, each grid point was colored to show the clade with the highest value, and the separation (self-organization) by clade is more clearly shown.

**Figure 1.**
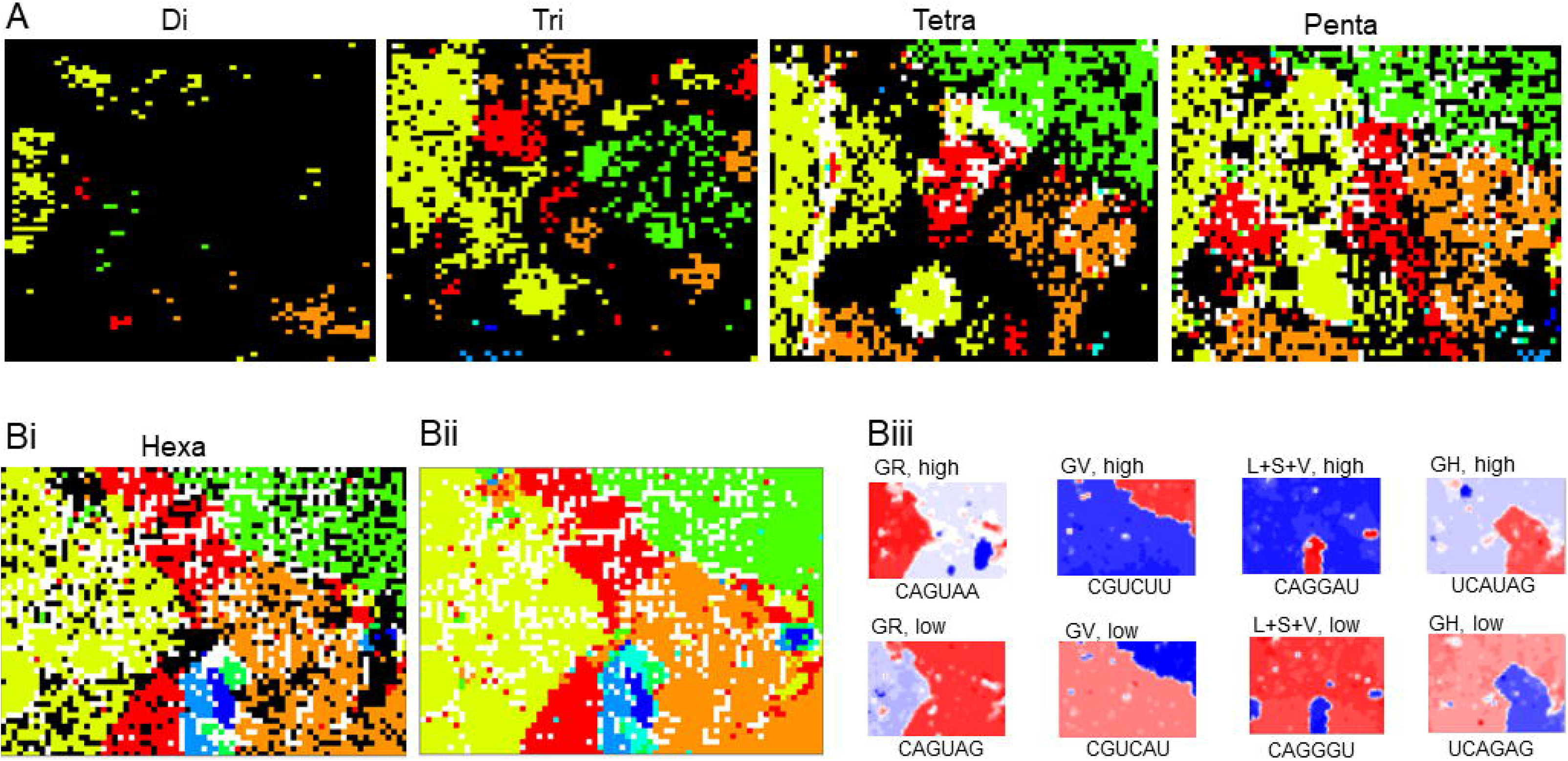
BLSOMs for di- to hexanucleotide compositions in SARS-CoV-2 genomes. **(A)** BLSOMs were constructed for di- to pentanucleotide compositions in 655,802 viral genome sequences. The total number of grid points (nodes) was set to 1/100 of the total number of genomes. Grid points that include sequences from more than one clade are indicated in black, and those containing sequences from a single clade are indicated in a clade-specific color: 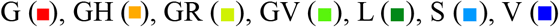 and 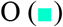. **(B)** BLSOMs for hexanucleotide composition. (i) Each grid point was displayed as described in A. (ii) Each grid point was colored for showing the clade with the highest value. (iii) Heatmaps for four pairs of hexanucleotides with one-base difference. More examples are presented in Supplementary Fig. S1.

### Explainable AI

The BLSOM is an explainable AI that can identify diagnostic oligonucleotides responsible for the observed clustering through the use of heat maps (Abe et al., 2003, 2006c). Eight examples of heatmaps for a hexanucleotide BLSOM are shown in Fig. 1Biii; here, the representative vector for each grid point is composed of 4,096 (= 4^6^) variables, and the contribution level of each variable at each grid point can be visualized as follows: high (red), moderate (white) and low (blue). For example, CAGUA<u>A</u> has a high occurrence (red) primarily in GR territories (yellow green in Fig. 1B), but CAGUA<u>G</u>, which differs from the former by one base (underlined), has a low occurrence (blue). When considering all 6-mer patterns (refer to Supplementary Fig. S1), we observed many cases in which red/blue patterns were reversed for a pair of 6-mers with a one-base difference, as observed in Fig. 1Biii.

It should be mentioned that most 6-mers exist in multiple copies in the viral genomes; thus, the one-base difference yielding the red/blue reverse pattern could not be connected uniquely to a mutation in the viral genome. In other words, extending the oligonucleotide length until most k-mers are present as one copy per genome should connect one-base difference to one mutation. Thus, we calculated occurrences of long oligonucleotides in the viral genomes and found that when the length was extended to 15-mers, most showed one copy per genome; i.e., if a 15-mer BLSOM yields a clade-specific separation (self-organization), this explainable AI will identify the mutations involved in the separation (Wada et al., 2020b; Ikemura et al., 2020). The 15-mers, however, include over one billion types (4^15^), and over 0.3 million types were found even for those that appeared in the viral genomes. A strategy for dimension reduction is essential for efficient data mining by a BLSOM.

### Time-series analysis for dimension reduction

A major internal structure, such as a clade, has arisen as a result of mutations that have occurred after the start of the epidemic and rapidly increased their population frequency. To study the formation of such internal structures using a 15-mer BLSOM, 15-mers in which no mutations have been found or those in which mutations have occurred but did not significantly expand in the virus population can be excluded from the analysis. Therefore, we first tabulated viral strains from each month of collection and calculated the occurrence of each 15-mer in the viral population. For each month, we calculated the increase in occurrence from December 2019 and selected 15-mers whose increase reached at least 10% in the population of each month; the reason for using the frequency of each month was to obtain information at intermediate stages of the pandemic (Wada et al., 2020b). After combining the results for each month and excluding duplicates, 891 different 15-mers were obtained. The BLSOM can analyze far more variables with no difficulty. Because various European-originated variants have shown worldwide spreads, the above time-series analysis was also performed for European strains, and 118 new 15-mers were obtained. When these are added, we may perceive the near-future trends of the global epidemic more acutely, and the following BLSOM was made using the 1009 15-mers (Fig. 2Ai); notably, the BLSOM can be used only for oligonucleotides of interest.

**Figure 2.**
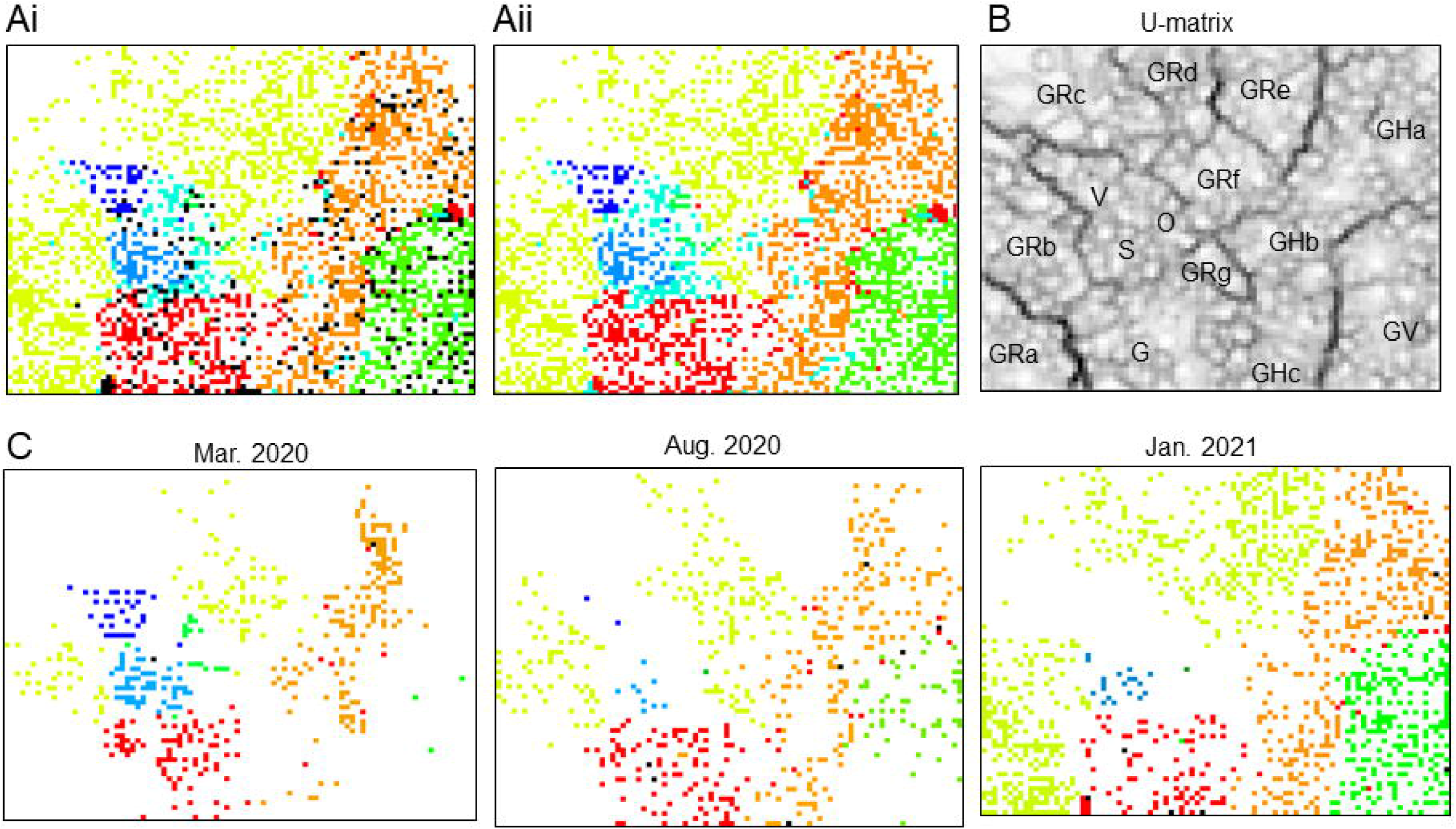
BLSOM for 1009 15-mers. The number of grid points was set as described in Fig. 1. **(A)** (i and ii) Each grid point was displayed as described in Fig. 1A and Bii, respectively. **(B)** U-matrix. The subgroupings in GR follow the dark black lines of U-matrix, while the others are primarily determined by heatmaps presented in Supplementary Fig. S2. (C) Sequences of three different collection months were displayed, and each grid point was marked as described in Fig. 1A.

### BLSOM for 15-mers

In Fig. 2Ai, each grid point of the BLSOM for the 1009 15-mers has 100 genomes on average, as above-mentioned in Fig. 1. Grid points that contain even one genome from another clade are shown in black, but most are colored, which shows that the BLSOM achieves good separation by clade, even though the number of oligonucleotides has been reduced by approximately a quarter compared to that obtained with the hexanucleotide BLSOM (4096 variables): usefulness of dimension reduction. Nevertheless, it is also clear that a significant portion was marked in black. Investigation of these black points revealed that they frequently contained the O (Other) clade sequences that GISAID did not classify as known specific clades. We believe that significant numbers of O-clade sequences can be classified into known clades (Ikemura et al., 2020). In Fig. 2Aii, each grid point was colored to indicate the clade with the highest value.

Based on the U-matrix (Fig. 2B) used in the conventional SOM analysis (Ultsch, 1993), the Euclidean distance between representative vectors of neighboring grid points can be visualized by the degree of blackness, and a larger distance is reflected by a higher degree of blackness (Iwasaki et al., 2013c). Notably, the boundary of the clade territory is visualized by a clear black line, and clear black lines are also observed inside the GR clade territory, indicating the existence of its subgrouping; the GR territory is very large, which reflects that GR sequences account for more than 60% of the total sequences. To clarify whether these GR subgroups have biological significance, we next investigated the time-series changes as follows. In Fig. 2C, using the BLSOM shown in Fig. 2A, sequences of different collection months are displayed. The comparison of March 2020, when almost all clades first appeared, with January 2021 showed that their sequences presented differential locations, and August 2020 showed an intermediate pattern. The existence of such time-series changes shows that the separation on the BLSOM has biological significance, as previously reported by Abe et al. (2021).

### Mutations responsible for separation into clades and their subgroups

We subsequently examined the 15-mers that contributed to separation on the BLSOM shown in Fig. 2 using the heatmap method explained in Fig. 1Biii. Importantly, we could identify mutations responsible for the separation into clades and their subgroups because all 15-mers that have rapidly expanded have one copy in the genome. Because the 1009 patterns include many similar patterns and are very complex as shown in Supplementary Fig. S2, efficient data mining appears to be difficult. Therefore, we introduce the BLSOM for the 1009 heatmaps. As a typical AI image processing, each heatmap can be converted to vectorial data of the dimension corresponding to the number of pixels, which was 6570 (approximately 1% of the total number of genomes) in the present analysis. When we created a BLSOM for the vectorial data of the heatmaps (abbreviated heatmap BLSOM), most grid points are black (Fig. 3A), which indicates that many heatmap patterns are very similar to each other. An examination of the black grid points revealed that one grid point was often composed of heatmaps of 15 different 15-mers, which were shifted by one base and added up to a 29-mer in total. A homology search (blastn) of the 29-mer against the standard SARS-CoV-2 sequence (NC_045512) showed a mutation in the center of the 29-mer, i.e., we could identify the involved mutations. When referring to all grid points on the heatmap BLSOM, the number of the attributed sequences was mainly a multiple of 15 or close to it. We subsequently explain actual cases in more detail.

**Figure 3.**
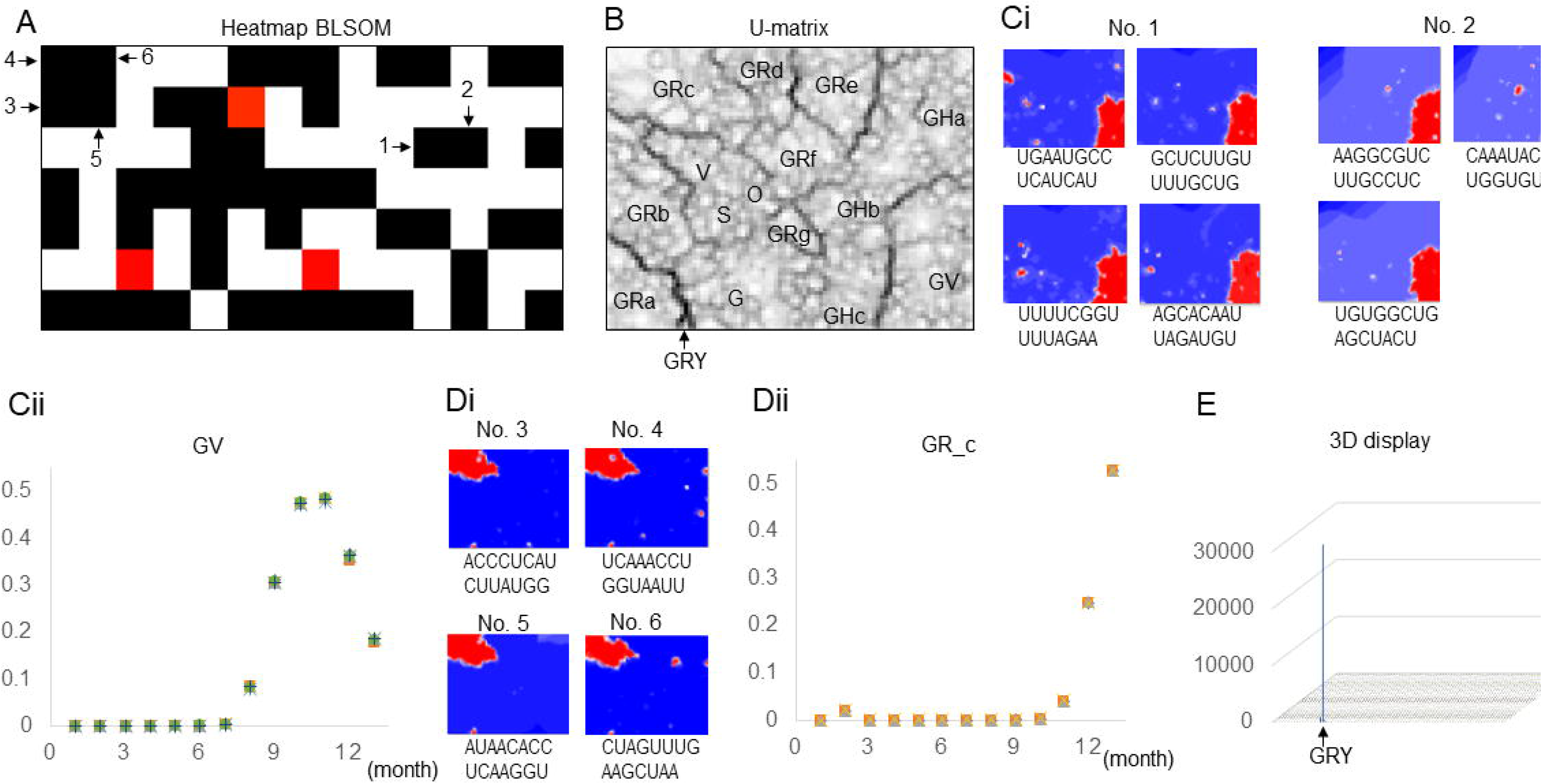
15-mers responsible for the separation according to clades and their subgroups. **(A)** BLSOM of heatmaps for 15-mer BLSOM presented in Fig. 2. Grid points (nodes) that contain one 15-mer heatmap are colored in red, and those including more than one heatmap are indicated in black. Four Grid points in focus are numbered. **(B)** The same to the U-matrix shown in Figure 2B, but the small GRY territory is marked with an arrow. **(C)** 15-mers contributed to formation of the GV territory: their heatmaps and monthly time-series changes (i and ii, respectively). Results of 15-mer sequences after mutation, which are listed in Table 1, are presented. In the time-series analysis, different 15-mers gave very similar patterns; thus, the symbols distinguishing individual 15-mers were not explained. **(D)** 15-mers contributed to formation of the GRc territory. Their heatmaps and time-series changes (i and ii, respectively) are shown as described in (C). **(E)** Mapping of newly downloaded GRY sequences on the 15-mer BLSOM presented in Fig. 2. The number of sequences mapped to each grid point is indicated by the height of the vertical bar: 3D display.

For the No.1 grid point in Fig. 3A, heatmaps of 60 (= 15 × 4) 15-mers were found; thus, four 29-mers were obtained from these 60 15-mers, as above-mentioned. The blastn analysis of these four 29-mers against the standard viral sequence identified four independent mutations. Table 1 shows the 29-mer sequences before and after the mutation and details of the mutation, as well as the four 15-mer sequences with the mutation at the center, for which actual heatmaps are presented in Fig. 3Ci. Although minor differences were found in the periphery of the GV territory, 15-mers with these four different mutations show a highly similar pattern (No. 1 in Fig. 3Ci). Fig. 3B shows the aforementioned U-matrix again. In addition to the GV territory, there are small red spots, which show differences among the four mutations. Importantly, over 0.6 million genomes were separated on a single BLSOM; thus, if a region of interest is found (e.g., red spots outside the GV territory), features of the corresponding genomes (e.g., the date and region of the strain collection) can be immediately displayed because this AI is equipped with various tools to enable efficient data mining.

For the No. 2 grid point in Fig. 3A, 45 (= 15 × 3) 15-mers were found, and their heatmap patterns (No. 2 in Fig. 3Ci) are very similar to those of No. 1, although small areas were missing in the GV territory. The resultant three 29-mers and corresponding mutations identified by blastn are also shown in Table 1. Figure 3Cii shows a monthly time-series analysis of a total of seven 15-mers presented in Fig. 3Ci, where 0 on the horizontal axis corresponds to December 2019 (the epidemic start) and other numbers show the elapsed months since the start. Their occurrences have increased worldwide since August 2020, until they reached a peak in November and decreased thereafter. Since these seven 15-mers show very similar time-series changes, the seven mutations of interest have increased and decreased, linking to each other. These mutations should include both advantageous mutations that increase infectivity and/or proliferation and neutral mutations that hitchhike the former mutations (Ikemura et al., 2020). The exploration of the seven mutations in Table 1 identified two nonsynonymous mutations and five synonymous mutations. The three mutations of No. 2 in Fig. 3Cii, where the red region is slightly missing, are synonymous mutations; thus, these may be neutral mutations. An AI that can visualize all genomes on a single map can support this type of prediction.

In the case of four grid points (No. 3 ~ 6 in Fig. 3A), 15 15-mers were attributed to each point, resulting in four 29-mers in total (Table 1). For four respective 15-mers in Table 1, most areas of GRc (a subgroup of GR) are red, but there are some differences in their periphery and spots in other clade territories (Fig. 3Di). Their time-series changes (Fig. 3Dii) are very similar to each other but different from those shown in Fig. 3Cii. As Figs. 3Ci and Di, we have introduced examples of heatmaps with relatively simple patterns among those in Supplementary Fig. S2 and identified mutations that expanded rapidly in the population. More analyses on the mutations have already been published (Wada et al., 2020b; Ikemura et al., 2020). Undoubtedly, the analysis aiming to distinguish between advantageous and neutral mutations becomes important, and the present AI is thought to be useful for this purpose. Extensive data of mutations observed for SARS-CoV-2 have been published by various sources, e.g., Alam et al. (2021) and COG-UK / Mutation Explorer (http://http://sars2.cvr.gla.ac.uk/cog-uk/).

### Use of preformed BLSOMs for analyses of newly-obtained sequences

One distinctive characteristic of SARS-CoV-2 is the rapid increase in sequence data, and GISAID has very recently added a new clade, GRY, which corresponds to variants originating in the UK and rapidly expanding worldwide. We have examined how the sequences belonging to the new clade are related to those analyzed in Fig. 2A. In Fig. 3E, the GRY sequences newly downloaded at the end of March 2021 (over 30,000 sequences), most of which were not included in the BLSOM in Fig. 2A, were mapped onto the BLSOM, i.e., for each GRY sequence, we looked for the grid point with the closest Euclidean distance. Most GRY sequences were mapped to the small area indicated by the arrow in Fig. 3B (arrowed also in Fig. 3E); here, the number of GRY sequences attributed to each grid point is indicated by the height of the vertical bar (3D display), and almost all GRY sequences were mapped in the GRY territory arrowed in Fig. 3B. The sequences downloaded at the beginning of February 2021 contained some GRY sequences, and these formed the small area in Fig. 2A. A BLSOM can provide information on sequences that were not included in its formation, and this mapping method is useful in metagenome analyses described below. So far, we have explained how the present AI can analyze big sequence data by explaining various methods in detail. Next, in connection with these detailed methodological strategies, we introduce examples of data mining from various prokaryotic and eukaryotic sequences.

### Large-scale BLSOMs and their use in metagenome studies

The BLSOM algorithm is suitable for high-performance parallel computing, and large-scale analyses, such as those that analyze most of the genome sequences currently available, is possible when using high-performance computers (Abe et al., 2006a). In Fig. 4A, the tetranucleotide composition of over 5 million 5-kb fragments of almost all microorganisms (including viral sequences) available in INSDC (DDBJ, ENA/EBI and NCBI) was analyzed by a BLSOM (Abe et al., 2020). In INSDC, only one strand of complementary sequences is registered, and the strand is chosen rather arbitrarily in the registration of fragment sequences. When investigating general characteristics of genome sequences, differences between two complementary strands are usually not important. In Fig. 4A, we constructed a BLSOM after summing frequencies of a pair of complementary oligonucleotides (e.g., AAAA and TTTT) in each fragment (Abe et al. 2005); BLSOM for this degenerate set of a pair of complementary tetranucleotides is abbreviated as DegeTetra BLSOM. Most of the sequences were separated (self-organized) with high accuracy according to their phylogeny with no information concerning phylogeny during the machine learning process (unsupervised AI). When such a large-scale BLSOM is created, the phylogeny of the metagenomic sequence can be predicted by mapping the large number of metagenomic sequences on the preformed BLSOM.

**Figure 4.**
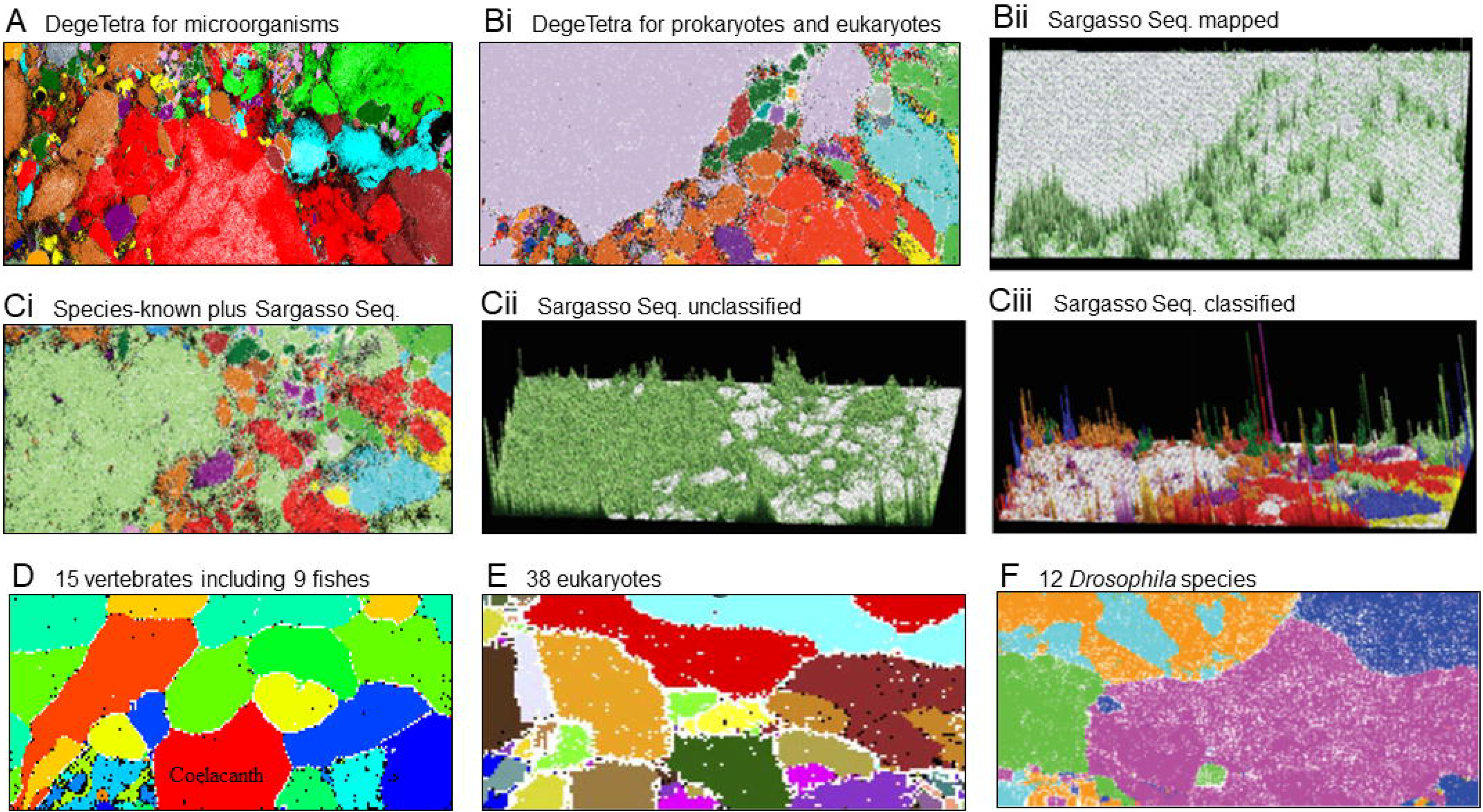
Oligonucleotide BLSOMs of various prokaryotic and eukaryotic genomes. **(A)** DegeTetra BLSOM of microorganisms of 40 phyla/classes; for details, see Abe et al. (2020). Grid points containing sequences of a single phylotype are indicated in phylotype-specific colors. **(B)** DegeTetra BLSOM of 211,000 and 210,600 5-kb sequences of 1,502 prokaryotes and 12 eukaryotes, respectively. (i) Grid points that contain sequences from a single prokaryotic phylotype are indicated in phylotype-specific colors, and those that contain only eukaryotic sequences are indicated in color 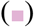; for details, see Abe et al. (2005). (ii) Metagenome sequences from Sargasso Sea (Venter et al., 2004) were mapped on this BLSOM, and square root of the number of Sargasso sequences mapped on each grid point is indicated by the height of the bar in color 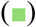. **(C)** DegeTetra BLSOM of species-known prokaryotic sequences and Sargasso sequences. BLSOM was constructed with 211,000 5-kb prokaryote sequences plus 218,000 Sargasso sequences after normalization of the sequence length; for details, see Abe et al. (2005). (i) Grid points that contain sequences from a single known phylotype are indicated in phylotype-specific colors, those that contain only Sargasso sequences are indicated in color 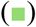, and those that include species-known and Sargasso sequences or those from more than one known phylotype are indicated in black. (ii) Square root of the number of Sargasso sequences classified into each grid point containing no species-known sequences is indicated by the height of the bar in color 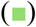. (iii) Square root of the number of Sargasso sequences that were classified into grid points containing species-known sequences is indicated by the height of the bar distinctively colored to show the phylotype. **(D)** DegeTetra BLSOM of 100-kb sequences of 15 vertebrates including nine fishes. For details, see Iwasaki et al. (2014). **(E)** DegeTetra BLSOM of 100-kb sequences of 38 eukaryotic genomes. For details, see Abe et al. (2006c). **(F)** DegePenta BLSOM of 100-kb sequences of 12 *Drosophila* species. After grouping the 12 species into five phylotypes, grid points are colored for showing the phylotypes; for details, see Abe et al. (2014).

We previously used this strategy to classify sequence fragments derived from environmental samples of the Sargasso Sea reported by Venter et al. (2004). Since a certain portion of Sargasso sequences may be derived from eukaryotic genomes, we constructed a DegeTetra BLSOM with 5-kb sequences from prokaryotes, unicellular eukaryotes and fishes (Fig. 4Bi); for details, see Abe et al. (2005). The power of BLSOM to separate prokaryotic from eukaryotic sequences was very high; only 0.1% prokaryotic sequences were classified into eukaryotic territories 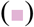. Next, we mapped Sargasso sequences longer than 1 kb on this BLSOM (Fig. 4Bii), as explained for the 3D display in Fig. 3E. While a major portion of the Sargasso sequences 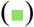) were classified into prokaryote territories, 9.9% were classified into eukaryotic territories that corresponded mainly to territories of unicellular eukaryotes.

This mapping method may have some disadvantages that even sequences derived from very novel phylotypes were attributed to any of known ones. To solve this issue, we next constructed a DegeTetra BLSOM with the species-known prokaryotic sequences plus Sargasso sequences (Fig. 4Ci). Grid points that contained only Sargasso sequences are indicated by 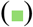, and the number of Sargasso sequences classified into these grid points is indicated by the height of the vertical bar (Fig. 4Cii). It is clear that a large portion of Sargasso sequences belongs to novel phylogenetic groups. In Fig. 4Ciii, the number of Sargasso sequences that got into grid points containing species-known sequences is indicated by the height of the bar distinctively colored to show the species-known phylotype (Abe et al., 2005).

The strategies explained in Figs. 4B and 4C have been used in various metagenome studies (Hayash et al., 2005; Uchiyama et al., 2005; Abe et al., 2006b; Uehara et al., 2011), including the detection of diverse pathogenic virus sequences from tick-derived metagenome sequences (Nakao et al., 2013). Dick et al. (2009) developed a slightly different type of SOM “ESOM (emergent SOM)” and obtained clear phylotype-specific classification of metagenomic sequences by analyzing several acidophilic biofilms in the Richmond mine.

### Analyses of various eukaryotic genomes

It is also interesting to see to what extent BLSOM can separate genome sequences of closely related species. Figure 4D shows DegeTetra BLSOM of 100-kb fragments from 15 vertebrates including nine fishes, and most sequences were separated by species with high accuracy (Iwasaki et al., 2014). Using heatmaps, such as explained in Figs. 1 and 3, we found that coelacanth has a distinctly lower frequency of CG-containing oligonucleotides than other fishes: an evident CG deficiency, as typically found for tetrapod. We also reported BLSOMs for 38 eukaryotes (Fig. 4E) (Abe et al. 2006c), 12 *Drosophila* species (Fig. 4F) (Abe et al., 2014), eight frogs (Katsura et al., 2021) and 13 plant species (Abe et al., 2009) and found high separation ability even for closely related species.

### Analyses of the human genome

Unsupervised AI can be used without prior knowledge or models, and we dare to leave the main data mining to the unsupervised explainable AI, which allows for unexpected knowledge discovery. Using pentanucleotide compositions of 50- and 100-kb and 1-Mb fragment sequences from the human genome, we explored the possibility of chromosome-dependent separation (Iwasaki et al., 2013c; Wada et al., 2015, 2020a). Although we first thought it was impossible, we obtained interesting patterns for these three lengths. Figure 5Ai shows a 50-kb case; grid points that contained only sequences from a single chromosome are colored, and those that contain sequences from plural chromosomes are shown in black. Most grid points were black, but four relatively large areas, as well as many small areas, contained characteristic colored grid points, and interestingly, the four areas were found to be composed of sequences derived from centromeric and pericentromeric heterochromatin regions (Iwasaki et al., 2013c); in the case of many small regions, sequences derived from subtelomeric heterochromatin regions were often found.

**Figure 5.**
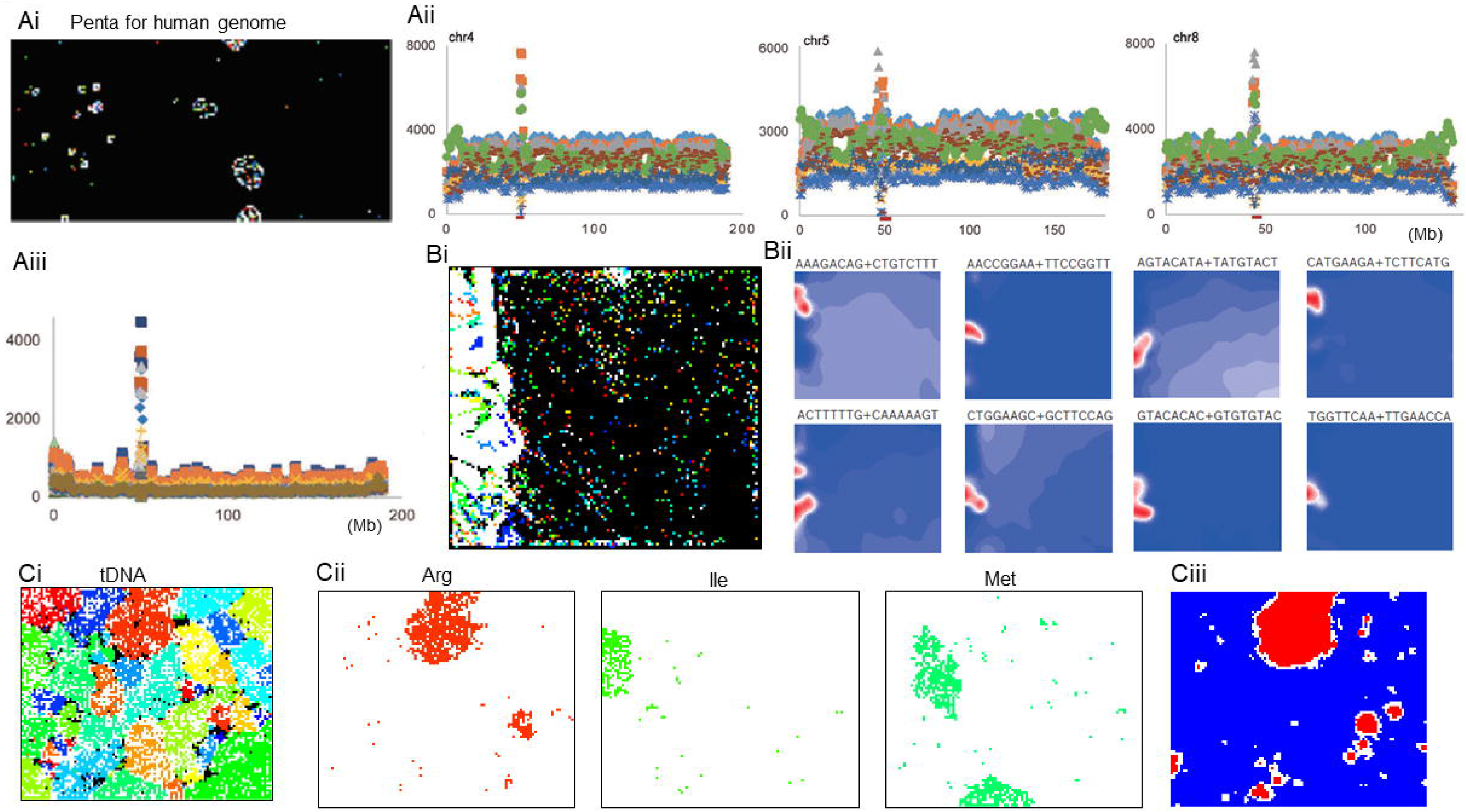
Analyses of the human genome and tDNAs. **(A)** Pentanucleotide BLSOM of 50-kb sequences derived from the human genome. (i) Grid points containing sequences from more than one chromosome are indicated in black, and those containing sequences from a single chromosome are indicated in chromosome-specific colors; for details, see Iwasaki et al. (2013c). (ii) Occurrences of eight TFBS-core pentanucleotides per Mb are plotted on three human chromosomes; a pair of complementary pentanucleotides were summed. Genomic regions of centromeric and pericentromeric constitutive heterochromatin are marked with a brown bar just below the horizontal axis; for details, see Iwasaki et al. (2013c). **(B)** BLSOM for 3946 octanucleotide TFBSs. A pair of complementary octanucleotide were summed. (i) Occurrences in 1-Mb sequences sliding with a 50-kb step were analyzes; for details, see Wada et al. (2020a). Grid points are marked as described in Ai. (ii) Eight examples of heatmaps to show the contribution level of each TFBS in each node. (iii) Occurrences of CG-containing pentanucleotides per 1 Mb are plotted as described in Aii. **(C)** Pentanucleotide BLSOM for bacterial tDNAs. (i) Grid points containing tDNAs of multiple amino acids are indicated in black, and those containing tDNAs of a single amino acid are colored; for details, see Iwasaki et al. (2017). (ii) Grid points containing tDNAs of individual amino acids on the Penta BLSOM are visualized separately with the color used there; 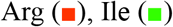 and 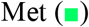. (iii) An example of heatmaps of diagnostic pentanucleotides responsible for amino-acid dependent clustering: ATAGA for Arg.

A heatmap analysis of the pentanucleotide BLSOM showed that the four large characteristic areas were evidently enriched in eight pentanucleotides that are known to be consensus-core sequences of diverse transcription factor-binding sequences (TFBSs) (Iwasaki et al., 2013c). Once the AI provides this information, we can analyze it more directly using a standard distribution map along each chromosome. Figure 5Aii displays the occurrences of the eight TFBS-core pentanucleotides in three chromosomes, and the TFBS-cores are clearly enriched in centromeric and pericentromeric regions, which are marked with a broad brown bar just below the horizontal axis. Importantly, this characteristic distribution pattern was observed in all chromosomes except chrY (Iwasaki et al., 2013c; Wada et al., 2020a).

For more direct analysis, we subsequently analyzed actual TFBSs, instead of the consensus-core pentanucleotides. By searching for human TFBSs through Swiss Regulon Portal (Pachkov et al. 2013), 189 hexanucleotide, 977 heptanucleotide and 3,946 octanucleotide TFBSs were found. When analyzing hexanucleotide TFBSs, their clear enrichment in centromeric and pericentromeric regions was observed as in Fig. 5Aii, whereas the combination of the enriched TFBSs varied among chromosomes; for details, see Wada et al. (2020a). Because the size of these enriched regions was at the Mb-level, we named the regions “Mb-level TFBS islands”. In centromeric and pericentromeric regions, the Mb-level enrichment in a group of CG-containing oligonucleotides (e.g., CG surrounded by A/T-rich sequences) was also observed (Fig. 4Dii); thus, these were named “Mb-level CpG islands”; actual sequences of the enriched oligonucleotides also varied among chromosomes (Wada et al., 2015).

### BLSOMs with TFBSs

In the case of 977 hepta- and 3,946 octanucleotide TFBSs, the number of variables is so large that it is practically impossible to perform a distribution map analysis of all TFBSs along all chromosomes. BLSOM, however, can analyze compositions even of 3,946 octanucleotide TFBSs in all chromosomes at once (Wada et al., 2020a). Figure 5Bi shows a BLSOM that analyzes the octanucleotide TFBS occurrences in 1-Mb sliding sequences with a 50-kb step. By restricting the oligonucleotides to TFBSs alone, characteristics of centromeric and pericentromeric regions became more prominent, and the sequences in these regions formed a large special area on the left side. Due to the 50-kb sliding step, the total number of 1-Mb sequences was almost the same to that in Fig. 5Ai where 50-kb fragments were analyzed, but the patterns were very different from each other for the following reasons. The TFBS occurrences in neighboring 1-Mb sequences off by 50 kb are very similar to each other, and therefore, sequences belonging to one chromosome can be visualized as a single stroke-like trajectory. Actually, in the large specific area, 1-Mb sequences derived from centromeric and pericentromeric regions of one chromosome give the colored dotted lines. Eight examples of heatmaps show chromosome-dependent enrichment in individual TFBSs (Fig. 5Bii), which is related in part to the chromosome-dependent difference in alpha-satellite monomer sequences (Hayden et al., 2013; Aldrup-MacDonald et al., 2016; Sullivan et al., 2017). Centromeric and pericentromeric regions are poor in protein-coding genes; thus, the evident enrichment in TFBSs was unexpected. In the previous paper (Wada et al., 2020), we proposed a model that their enrichment in the Mb-level structures (Mb-level TFBS islands) support interchromosomal interactions in interphase nuclei (Lieberman-Aiden et al., 2009; Dekker & Misteli, 2015).

### Analyses of tRNAs, proteins and codon usage

We will next explain what a BLSOM can tell us about short sequences such as tRNAs. Even if the result is unpredictable, we dare to leave knowledge discovery to AI. We have created a database (tRNADB-CE) of tRNA genes (tDNAs) obtained from all microbial genomes available (Abe et al., 2011, 2014), and we next explain a pentanucleotide BLSOM of approximately 0.4 million tDNA sequences in the database (Iwasaki et al., 2017). There are 1024 types of pentanucleotides, and sequences of tDNAs are clearly shorter than 100 nt; thus, most pentanucleotides do not occur in each tDNA. For these particular data consisting mostly of zeros, the obtained result was rather unpredictable. After training only with the pentanucleotide compositions, the grid points that contained tDNAs of one amino acid were colored. Interestingly, most tDNAs were separated by amino acids with high accuracy (Fig. 5Ci). Referring to heatmaps (Fig. 5Ciii), the oligonucleotides that contributed to the separation by amino acid were found to correspond primarily to the recognition sequences of the aminoacyl-tRNA transferase, which have been designated identifier or identity (Ardell, 2010), as well as the anticodon and surrounding nucleotides, i.e., the identifiers appear to be assignable without experiments (Iwasaki et al., 2017). Interestingly, a minor proportion of tDNAs were located outside the main territory of each amino acid on the BLSOM (Fig. 5Cii), and these exceptional tDNAs often belonged to phylogenetic groups that grow in special environments (Iwasaki et al., 2017). The BLSOM can provide insight into identifiers even for these poorly characterized species. An analysis of large numbers of tDNAs from a wide range of phylotypes will enable this type of characteristic research.

As introduced in Fig. 3A, the BLSOM can analyze vectorial data other than oligonucleotide compositions, and the peptide composition of proteins is analyzable. In fact, Ferran et al. (1994) investigated di- and tripeptide compositions using the Kohonen conventional SOM. Although the study was conducted at a time when a limited number of protein sequences (approximately 2000) were known, the proteins were separated by function. In addition, rather than using 20 different amino acids separately, 11 groups, into which amino acids with similar physicochemical properties were classified, achieved better separation. We created BLSOMs of di-, tri- and tetrapeptides for over 80,000 prokaryotic proteins and obtained separation reflecting COGs (Clusters of Orthologous Groups of proteins), which are known to be functional groups (Abe et al., 2009). In support of the previous research, a better separation was obtained after grouping into 11 groups. However, a mixed trend of separation reflecting phylogeny was detected, and the situation becomes further complicated for proteins with multiple functional domains. Although it has already been used to study the diversity in enzymes related to secondary metabolic pathways in plants (Ikeda et al., 2013), the scope of its use is currently limited, and further development is needed. BLSOM can also be applied to the studies of codon usage in protein genes (Kanaya et al., 2003; Kosaka et al., 2008).

## CONCLUSION AND FUTURE PERSPECTIVES

In genetics and related fields, most studies have been conducted by creating models and hypotheses and then testing them. However, in the big data era, this principle may not be sufficient for the data mining of big data. Models and hypotheses must rely on existing knowledge, which make it difficult to obtain unexpected insights. Here, we have explained the strategy of entrusting knowledge discovery to AI, which appears to become increasingly important. In big data analyses, the research conducted by large groups using high-performance computers is becoming the norm, and somewhat unfortunately, the contributions of individuals and small groups are decreasing. With the help of AI, individuals and small groups can discover distinctive knowledge from big data. The research presented here, with the exception of those shown in Figs. 4A~C, can be conducted on PC-level computers. A BLSOM program suitable for PC systems is available on the following website: http://bioinfo.ie.niigata-u.ac.jp/?BLSOM.

The main purpose of this paper was to show how an unsupervised explainable AI can enable knowledge discovery from big data. To help readers envision applications to their own topics, various examples and research strategies were explained in detail. The present research strategies can be applied to AI algorithms other than the BLSOM, and similar or better data mining will become possible. The Kohonen SOM, which serves as the basis of the BLSOM, was developed during the second AI boom and is a classic technology. In the current third AI boom, a wide variety of new technologies are being developed and playing active roles in practical fields. For an example of image processing, image classification has made great strides due to its practical importance, but the reasons for why AI is achieving such highly accurate classification are unclear for us (black box). This may not be a problem regarding the practical use, but in scientific studies, the reason for the classification is important. The third-generation explainable AI will be definitely developed. On the machine side, quantum computers, which excel at combinatorial optimization problems, have become practical. The BLSOM algorithm appears to somewhat resemble the combinatorial optimization problem; hence, AI-based genome analyses using quantum computers will be realized sooner than expected. Such advanced technologies should be quickly introduced to the studies with social importance, such as observed for the rapid technological development of COV-19 vaccine production. In such an era, interdisciplinary collaborative research and appropriate education will become increasingly important.

## ACKNOWLEDGEMENTS

We gratefully acknowledge the authors publishing their sequences from GISAID’s Database. This work was supported by JSPS KAKENHI Grant Number 18K07151, JST CREST Grant Number JPMJCR20H1 and AMED under Grant Number JP20he0622033.

## Notes

### Competing Interest Statement

The authors have declared no competing interest.

