## Supplementary Figs S1 and2 for "Unsupervised explainable AI for the collective analysis of a massive number of genome sequences: various examples from the small genome of pandemic SARS-CoV-2 to the human genome"

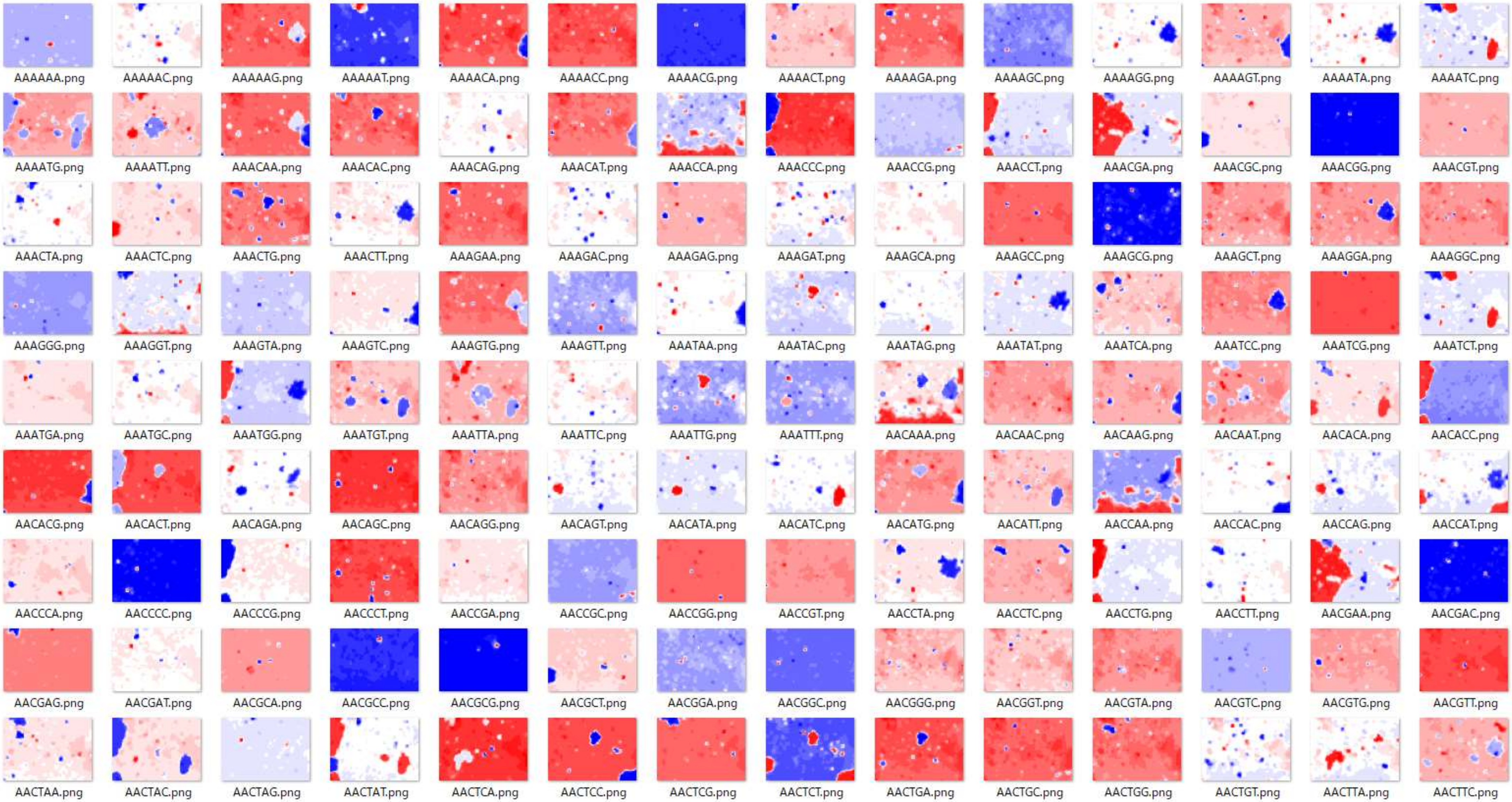


Supplementary Fig. S1. A portion of heatmaps of hexa BLSOM. The “T” base, instead of “U”, was used for compatibility with the NCBI reference genome.

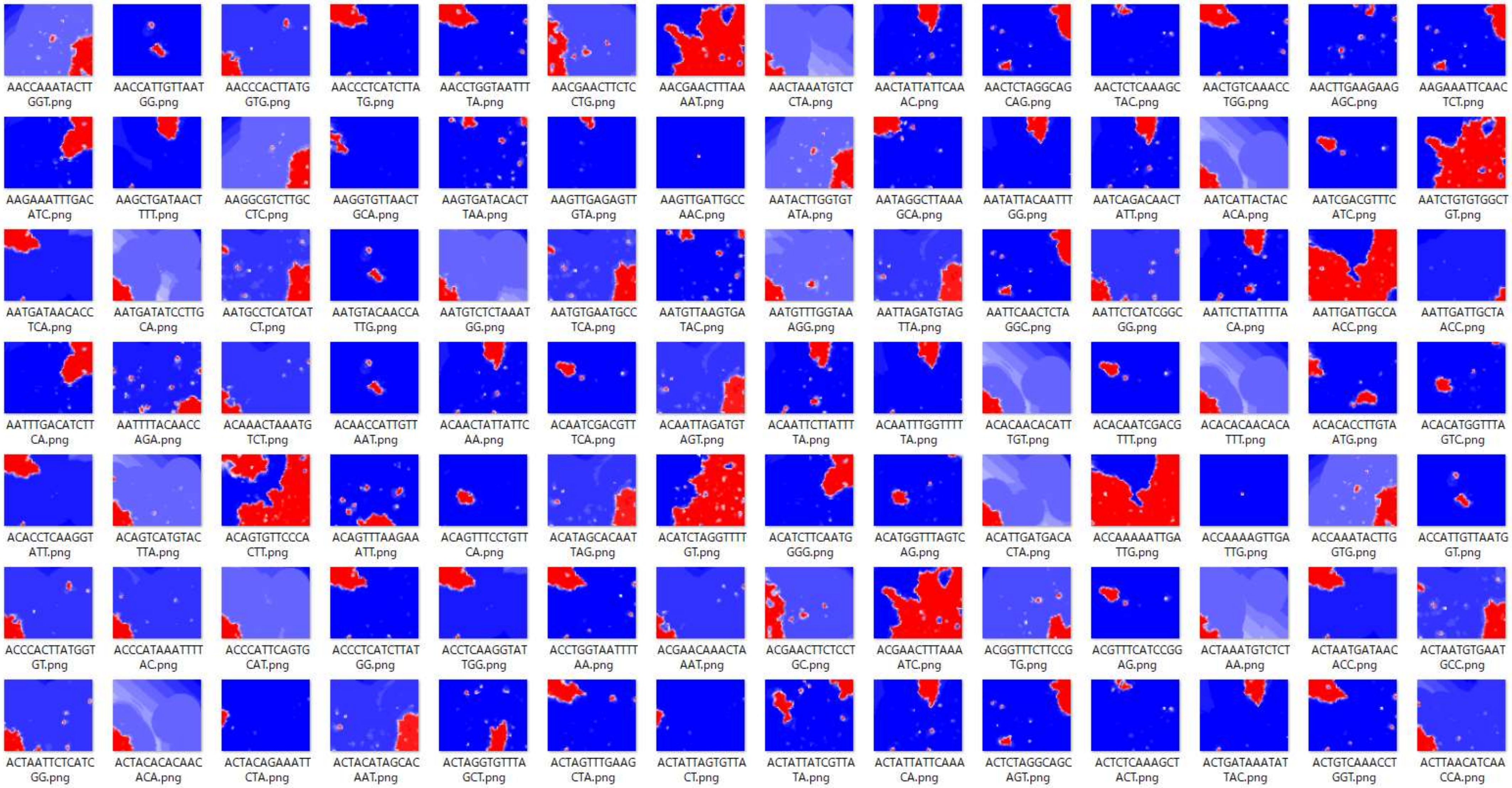


Supplementary Fig. S2. A portion of heatmaps of 15-mer BLSOM. The “T” base, instead of “U”, was used for compatibility with the NCBI reference genome.
